# Recognition of pathogenic bacteria by intestinal progenitors promotes adult *Drosophila* midgut regeneration via PGRP-MKK3-p38 signalling

**DOI:** 10.1101/2025.05.22.655543

**Authors:** Bhavin Uttekar, Rebecca Wafer, Suci Cendanawati, Parthive H. Patel

**Affiliations:** School of Cellular and Molecular Medicine, Faculty of Health and Life Sciences, University of Bristol, University Walk, Bristol BS8 1TD UK

## Abstract

When enteropathogenic bacteria breach the intestinal epithelium, they are recognized by epithelial and immune cells that elicit an intestinal regenerative response. However, less is known about whether and how intestinal progenitors directly detect invading pathogenic bacteria and couple this to their proliferation. Here we show that adult *Drosophila* midgut progenitors recognise peptidoglycan from pathogenic bacteria through the peptidoglycan recognition proteins, PGRP-LC and PGRP-LE, and translate this to ISC proliferation by stimulating MKK3-p38 signalling. Moreover, we find that PGRP-LC/LE-MKK3-p38 signalling in progenitors regulates p38 activation throughout the midgut epithelium after infection, indicating that progenitors can influence the regenerative niche in a non-cell autonomous manner. Whilst it was previously thought that ISC proliferation in both mammals and flies is driven solely by damage-induced signals after infection, our work reveals that progenitors can directly recognise pathogenic bacteria and mount a strong parallel regenerative response that spreads throughout the midgut epithelium. Increased ISC proliferation after bacterial recognition may also serve as a strategy to repopulate the epithelium with uninfected cells.

## Introduction

The inability to properly maintain or regenerate an epithelium leads to a breakdown of its integrity, structure and function, ultimately disrupting organismal homeostasis ^1, 2^. Maintenance of the mammalian intestinal epithelium relies on intestinal stem cells (ISCs), which generate diXerentiated absorptive and secretory epithelial cells ^3^. In response to intestinal damage, ISCs receive proliferative signals, including Epidermal Growth Factor (EGF), cytokines, and Wnts to replenish damaged or lost cells ^4, 5^. These signals originate from a regenerative niche composed of resident epithelial cells, immune cells, mesenchyme, vasculature, and enteric glial cells ^4, 5^.

Enteropathogenic bacterial infections cause intestinal damage. When enteropathogenic bacteria eventually breach the intestinal epithelial barrier, they are recognized by intestinal epithelial and innate immune cells ^6^. This results in cytokine production and inflammation that supports a regenerative response within the intestinal epithelium ^6^. While much is known about how intestinal epithelial and immune cells sense invading pathogenic bacteria, we know little about whether ISCs or intestinal progenitors sense bacteria during infection and mount a regenerative response.

Here we address this question using *Drosophila* as a model to study the response of midgut progenitors to bacterial intestinal infections. Like the mammalian intestine, the adult *Drosophila* midgut epithelium is maintained by ISCs that generate two primary epithelial cell types: absorptive enterocytes (ECs) and secretory enteroendocrine cells (EEs) ^7–9^. In approximately 90–95% of cases, ISCs divide to produce an enteroblast (EB)— the precursor for ECs ^8, 10, 11^. When EBs then receive a strong Notch signal from the ISC, they differentiate into ECs ^10, 12^. In contrast, EEs arise either from ISCs via an enteroendocrine precursor (EEp) or through direct differentiation of ISCs ^13^. ISCs and their daughters, EBs and EEps, are collectively considered adult midgut progenitors and all express the progenitor marker *escargot* (*esg*) ^7^. Together with the visceral muscle and trachea that surround the midgut epithelium, EBs, ECs, and EEs form a regenerative niche that facilitates midgut repair by producing signals that directly promote ISC proliferation ^8, 14, 15^. These cues include Unpaired cytokines (Upd1-3), EGFs, Wnt, Hedgehog, fibroblast growth factor (FGF), and PTK7/Off-track ^14–23^.

Oral infection of adult flies with gram-negative pathogenic bacteria such as *Pseudomonas entomophila* (*P.e.*) or *Erwinia carotovora carotovora 15 (Ecc15)* causes midgut damage ^19, 24–26^. In the case of *P.e*., this is due to the expression of virulence factors that directly cause midgut damage, especially the pore-forming toxin, Monalysin ^27, 28^. Damage to the midgut epithelium induces a regenerative niche that stimulates ISC proliferation ^19, 29^. Nevertheless, it is unknown whether bacterial recognition by midgut cell types plays a direct role in the regenerative response. Similarly, the human intestine can be infected by gram-negative pathogenic bacteria that damage epithelial cells, including *Escherichia coli*, *Shigella flexneri* and *Salmonella serovar* ^30^. While commensal bacterial recognition by ISCs has been suggested to promote intestinal homeostasis ^31, 32^, it is unknown whether our intestines, particularly the ISCs, sense invading pathogenic bacteria and mount a regenerative response.

In the adult *Drosophila* midgut, ECs can recognise invading pathogenic bacteria (*Ecc15*) by sensing peptidoglycan, a component of the bacterial cell wall, through peptidoglycan recognition proteins (PGRPs) ^33^. Peptidoglycan is a crosslinked polymer, with structural variations distinguishing gram-positive (Lys-type) and gram-negative (Dap-type) bacteria ^34–36^. *Drosophila* PGRPs are classified by size into short extracellular forms (PGRP-S) and long forms (PGRP-L) that can be extracellular, intracellular and transmembrane ^34, 36^. The long forms of PGRP (PGRP-LC and PGRP-LE) detect DAP-type peptidoglycan mainly from gram-negative bacteria and activate antimicrobial production (AMP) via the immune deficiency (IMD) pathway ^36–38^. Further, both PGRP-LC and PGRP-LE are important in ECs of the posterior adult midgut for AMP production after *Ecc15* infection ^37, 38^. Additionally, *Ecc15* peptidoglycan can activate p38 signalling in ECs through IMD-Mekk1-MKK3 signalling ^33^. p38 signalling in ECs promotes the expression of the NADPH oxidase, Dual oxidase, resulting in increased ROS production ^33^. This increased ROS production eliminates invading pathogenic bacteria from the lumen ^33, 39^. Together, peptidoglycan sensing by ECs elicits two strategies—ROS and AMP production—to eliminate invading pathogenic bacteria from the lumen. Although it was hypothesized that ROS production triggered by *Ecc15* peptidoglycan sensing in ECs indirectly causes epithelial damage and ISC proliferation, this was found to not be the case ^40^. Thus, it remains unknown whether midgut cell types, particularly ISCs, can recognise peptidoglycan and contribute directly to regeneration.

In mammals, most peptidoglycan sensing by cells likely occurs through intracellular Nod1 and Nod2 receptors ^35^. Their functional equivalent in *Drosophila* is PGRP-LE ^36, 37^. Nod1 senses -negative bacteria, whilst Nod2 senses gram-positive and gram-negative bacteria ^35^. Both Nod1 and Nod2 interact with the receptor-interacting serine/threonine protein kinase 2 (RKP2), activating either nuclear factor-Kß (NF-Kß), or mitogen-activated protein kinase kinase kinase 7 (TAK1/MAP3K7) and mitogen-activated protein kinase (JNK, p38) signalling to stimulate cytokine production ^35^. Nod2 has been shown to be expressed at higher levels in Lgr5^+^ stem cells within intestinal organoids. Treatment of intestinal organoids with the Nod2 ligand muramyl dipeptide—a component of bacterial peptidoglycan—produces a higher yield of intestinal organoids, suggesting that bacterial sensing could promote ISC survival and intestinal homeostasis ^31^. Further, Nod2 was found to be critical for the regeneration of ISCs and intestinal epithelial restitution following doxorubicin-induced ISC and crypt cell depletion ^31^, which does not occur upon infection with pathogenic bacteria. This study did not examine a pathophysiological in vivo role for Nod2 in intestinal regeneration during pathogenic bacterial infection, where ISCs are spared.

Here we explore if and how adult *Drosophila* midgut progenitors recognise pathogenic bacteria after infection and translate this into a regenerative response. We find that PGRP-LC/LE-MKK3-p38 signalling is required within progenitors for adult *Drosophila* midgut regeneration after pathogenic bacterial infection. Mechanistically, PGRP-LC and PGRP-LE in progenitors recognise peptidoglycan and translate this to ISC proliferation by activating MKK3-p38 signalling. Furthermore, we show that PGRP-LC/LE-MKK3-p38 signalling in progenitors activates p38 throughout the epithelium after infection, indicating that progenitors can shape the regenerative niche in a non-cell autonomous manner to support epithelial regeneration. While ISC proliferation after intestinal infection by pathogenic bacteria has been thought to be solely due to midgut damage, our work shows that ISC proliferation is also strongly induced by direct pathogen recognition by progenitors.

## Results

### p38 signalling promotes adult *Drosophila* ISC proliferation after pathogenic bacterial infection

Previously, it was shown that infection with pathogenic gram-negative bacteria—including *Ecc15* and *P.e*.—activates p38 signalling in ECs of the adult *Drosophila* midgut epithelium*. Ecc15* infection induces p38a in ECs through peptidoglycan recognition by PGRP-LC and subsequently via IMD-Mekk1-MKK3 signalling ^33^. After *P.e.* infection, however, p38a and p38b are activated in ECs through damage sensing by Nox-Ask1-MKK3 signalling ^41^. Using an antibody against phosphorylated human p38 (Thr180/Tyr182)(p-p38), we found increased levels of activated p38 within *esg^+^* progenitors in addition to ECs after *P.e.* infection compared to the basal p-p38 levels found in control progenitors (Figure 1A-C). This increased phosphorylated p38 likely detects both activated p38a and p38b within progenitors ^41^. p38 was induced by 134.8% within *esg^+^* progenitors after *P.e.* infection compared to uninfected midguts (Figure 1D).

**Figure 1.**
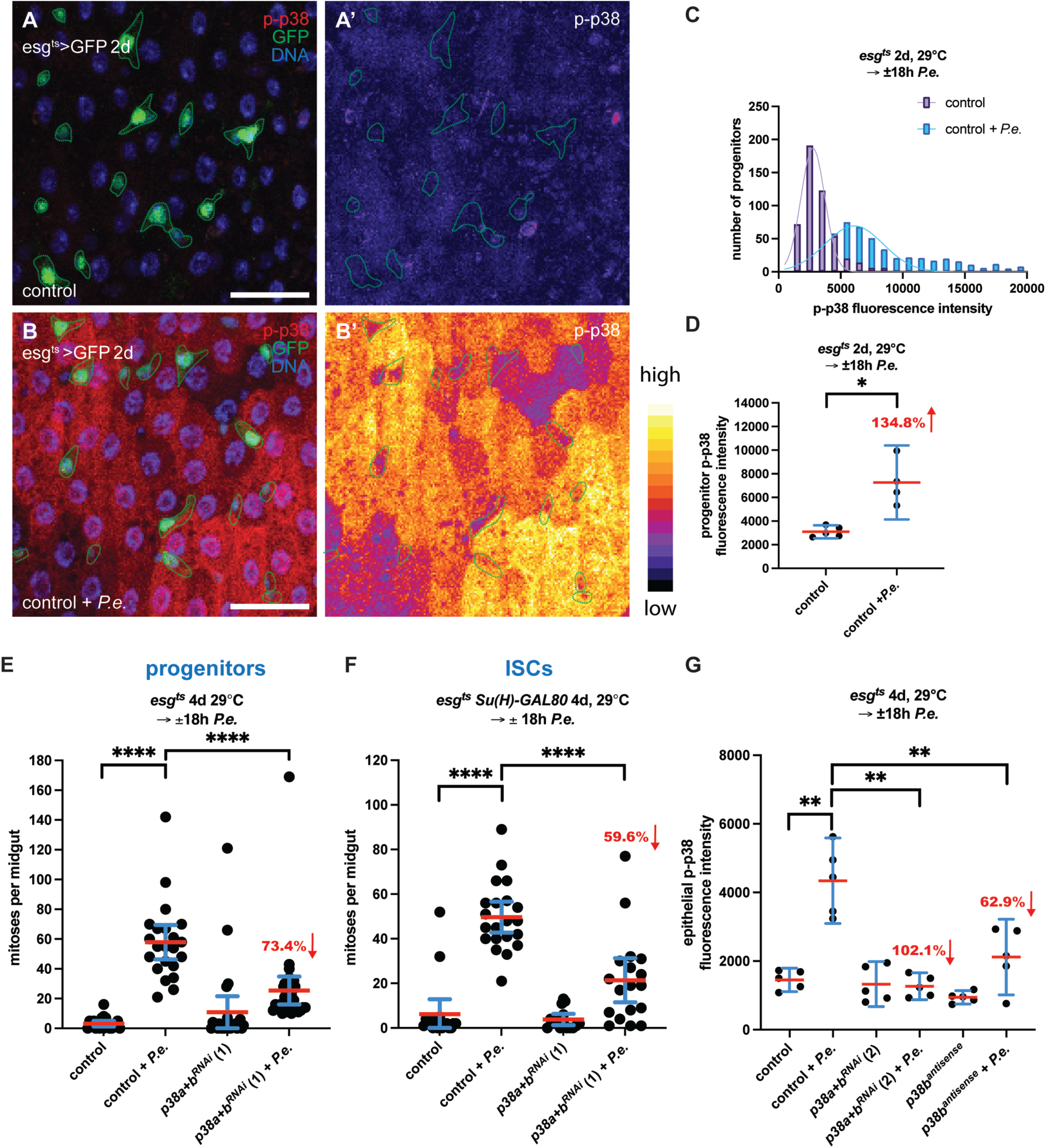
p38 activation within progenitors after pathogenic bacterial infection induces ISC proliferation and p38 activation throughout the adult *Drosophila* midgut epithelium. p38 is activated in progenitors after *P.e.* infection **(A-D)**. Phosphorylated-p38 (p-p38) (A, B: red; A’, B’: LUT) levels increase throughout the midgut epithelium including the progenitors (GFP in A, B; green-dashed outline in A-B’) in *P.e.* infected midguts compared to uninfected control midguts. The distribution (histogram) of p-p38 fluorescence intensity per progenitor from control uninfected midguts (n=5 midguts from one of two experiments) and *P.e.*-infected midguts (n=4 midguts from one of two experiments) (C). The mean p-p38 fluorescence intensity in progenitors per midgut with mean and 95% CI are shown for control midguts (n=5 from one of two experiments) and *P.e*.-infected control midguts (n = 4 from one of two experiments) (Mann–Whitney; p-value = 0.0159) (D). p38 is required in ISCs for their proliferation **(E-F)**. Depleting *p38a* and *p38b* in progenitors with *esg^ts^* and in ISCs with *esg^ts^Su(H)Gal80* for 4 days blocks ISC proliferation by 73.4% (Mann–Whitney; p < 0.0001) and 59.6% (Mann–Whitney; p < 0.0001), respectively, compared to *P.e.*-infected control midguts. The mean number of PH3-positive cells per midgut with 95% CI is shown for control midguts (n = 21 pooled from three experiments), *P.e.* infected control midguts (n = 22 pooled from three experiments), midguts expressing *p38a+b^RNAi^* (1) in progenitors for 4 days (n = 26 pooled from three experiments) and *P.e.*-infected guts expressing *p38a+b^RNAi^* (1) in progenitors for 4 days (n = 34 pooled from three experiments) (E). The mean number of PH3-positive cells per midgut with 95% CI in control midguts (n = 18 pooled from two experiments), *P.e.*-infected control midguts (n = 21 pooled from two experiments), midguts expressing *p38a+b^RNAi^* (1) in ISCs for 4 days (n = 15 pooled from two experiments) and *P.e.*-infected midguts expressing *p38a+b^RNAi^* (1) in ISCs for 4 days (n = 18 pooled from two experiments) (F). In E and F, the mean number of mitoses per midgut increased in *P.e.-*infected control midguts compared to uninfected midguts (Mann-Whitney; p<0.0001). p38 is required in progenitors for p38 activation throughout the midgut epithelium after *P.e.* infection **(G)**. p38 fluorescent intensity significantly increases throughout the midgut epithelium after *P.e.* infection. Depletion of either *p38a* and *p38b or p38b* alone in progenitors with *esg^ts^* for 4 days reduces p38 activation throughout the midgut epithelium after *P.e.* infection by 102.1% (Mann-Whitney; p value = 0.0079) and 62.9% (Mann-Whitney; p-value = 0.0079), respectively, compared to control infected midguts. The mean fluorescence intensity for epithelial p38 is shown for control midguts (n = 5 from one experiment), *P.e.*-infected control midguts (n = 5 from one experiment), midguts expressing *p38a+b^RNAi^* (2) in progenitors (n = 5 from one experiment) and *P.e.*-infected midguts expressing *p38a+b^RNAi^* (2) in progenitors (n = 5 from one experiment). The mean fluorescence intensity for epithelial p38 increased in *P.e.-*infected control midguts compared to uninfected midguts (Mann-Whitney; p-value = 0.0079) (G). Scale bars are 30μm.

Since p38 was activated within progenitors after *P.e.* infection, we next tested whether p38 signalling was required in progenitors for ISC proliferation after *P.e.* infection. To do this, we depleted p38 signalling from midgut progenitors by expressing an RNAi that targets both p38a and p38b (*p38a+b^RNAi^*) using the progenitor-specific driver *esg-GAL4 tubGAL80^ts^* (*esg^ts^*) and infected these flies with *P.e.* (Figure 1E). Depleting p38a and p38b in progenitors resulted in a 73.4% decrease in ISC proliferation after *P.e.* infection compared to infected control midguts. This decrease was not caused by reduced progenitor cell number due to *p38a and p38b* depletion with *esg^ts^* and *P.e.* infection as the mean percent *esg^+^* progenitors in these midguts was similar to uninfected and *P.e*- infected control midguts (Supplementary Figure 1A).

*esg^+^* midgut progenitors include ISCs and their daughters, EBs and EEps. Thus, we could not conclude from our data whether p38a and p38b were required in ISCs or in their daughters for their proliferation after *P.e.* infection. To determine this, we depleted *p38a* and *p38b* from ISCs using an ISC-specific driver, *esgGAL4 tubGAL80^ts^ Su(H)Gal80* (*esg^ts^ Su(H)Gal80*). This system allowed us to express *p38a+b^RNAi^* in ISCs whilst blocking GAL4- mediated expression in *Su(H)^+^* EBs by Gal80 expression. We found that p38a and p38b depletion in ISCs strongly blocked their proliferation after *P.e.* by 59.6% (Figure 1F) compared to control infected midguts. These data suggest that p38a and p38b signalling is required in ISCs to respond to midgut damage.

Lastly, we examined whether blocking p38 signalling in progenitors aXects their levels of activated p38 in progenitors after *P.e.* infection. We found that depletion of p38b resulted in a 72.1% decrease in p38 activation in progenitors after infection compared to control infected midguts (Supplementary Figure 1B-C). Surprisingly, we also found that blocking p38 activity specifically in progenitors decreased the levels of activated p38 throughout the midgut epithelium. Depletion of p38a and p38b in progenitors with *esg^ts^* blocked p38 activation after *P.e.* infection throughout the midgut epithelium by 102.1% (Figure 1G, Supplementary Figure 1D-G). Further, depletion of p38b alone in progenitors blocked p38 activation in the entire midgut epithelium by 62.9% (Figure 1G). These data indicate that in addition to regulating ISC proliferation, p38a and p38b activity in progenitors non-cell autonomously controls p38 activity throughout the midgut epithelium.

### MKK3-p38 signalling in intestinal progenitors induces ISC proliferation and p38 activation in the midgut epithelium

We next determined whether increasing p38 activity in midgut progenitors is suXicient to stimulate ISC proliferation. To activate p38 signalling within midgut progenitors, we overexpressed the *Drosophila* MKK3, Licorne, with *esg^ts^*. Overexpression of Licorne in progenitors caused a significant increase in ISC proliferation after 1 day of expression (Figure 2D). Furthermore, we found that Licorne overexpression in progenitors for 1 day with *esg^ts^* increased p38 activation by 215.2% within progenitors (Figure 2A-C, Supplementary Figure 2A). Interestingly, we also found increased p38 activation in ECs neighbouring Licorne-overexpressing progenitors (Figure 2A-B’). Together, these data show that overexpressing MKK3 in progenitors can promote ISC proliferation and p38 activity in both progenitors and surrounding ECs.

**Figure 2.**
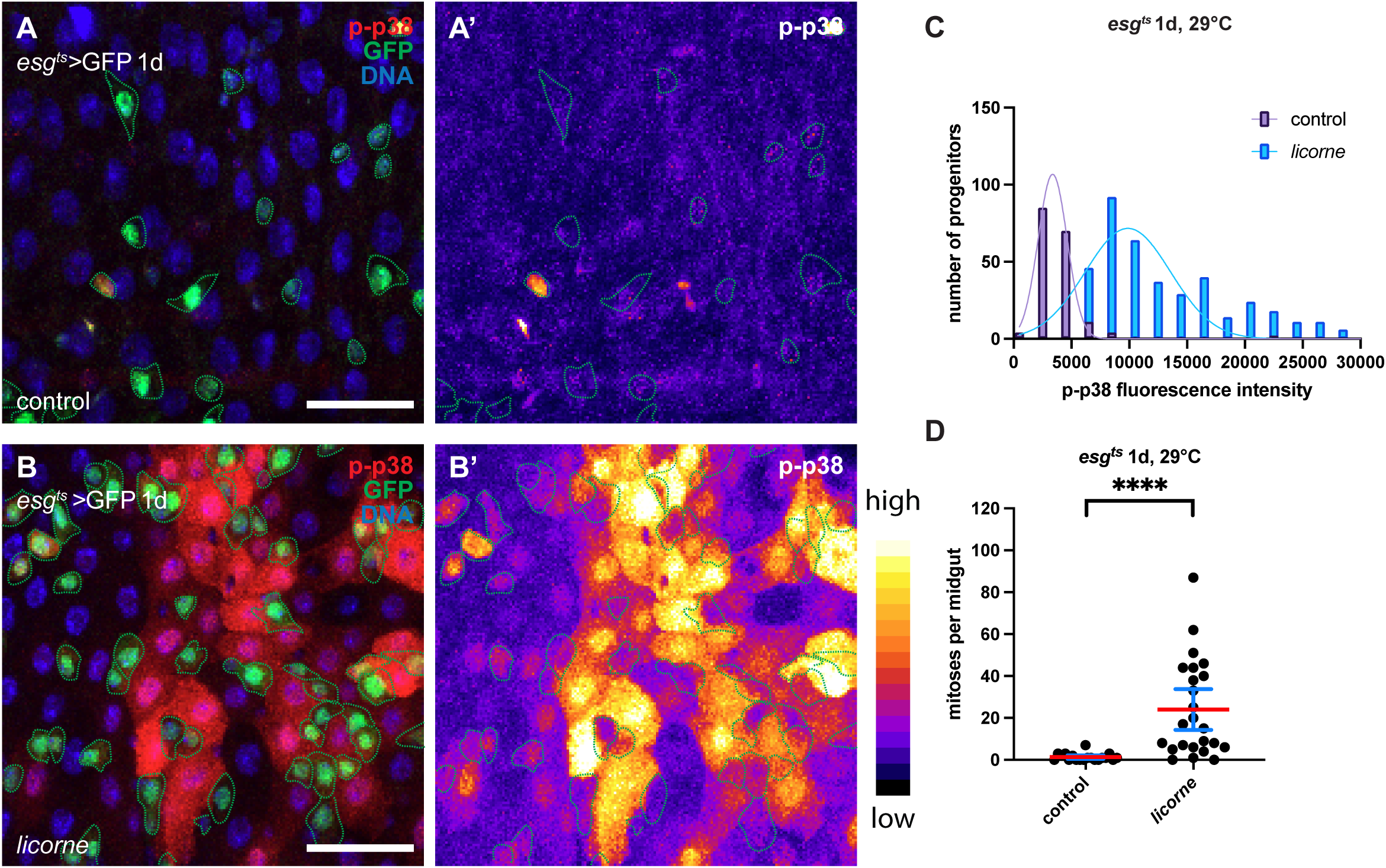
MKK3/Licorne overexpression in progenitors activates p38 within progenitors and stimulates intestinal stem cell proliferation. Licorne overexpression in progenitors activates p38 in progenitors and in nearby midgut epithelial cells **(A-C)**. Phosphorylated-p38 (p-p38) (A, B: red; A’, B’: LUT) increases in midgut progenitors (A, B: GFP; A-B’, green-dashed outline) and in nearby epithelial cells after overexpressing *licorne* with *esg^ts^* compared to control midguts. The distribution (histogram) of p-p38 fluorescence intensity per progenitor is shown for control midguts (n=5 from one experiment) and midguts overexpressing *licorne* in progenitors for 1 day with *esg^ts^* (n=5 from one experiment). Licorne overexpression in midguts promotes ISC proliferation **(C)**. The mean number of PH3-positive cells per midgut with 95% CI increases in midguts overexpressing *licorne* in progenitors for 1 day (n = 24 from one experiment) compared to control midguts (n = 19 from one experiment) (Mann–Whitney; p < 0.0001) (D). Scale bars are 30μm.

Since Licorne overexpression stimulated ISC proliferation in the absence of *P.e.* infection, we next tested whether Licorne was required for ISC proliferation after *P.e.* infection. We found that depleting *licorne* with two RNAis (RNAi(1) and RNAi(2)) in progenitors with *esg^ts^* blocked ISC proliferation after *P.e.* infection by 97.7% and 36.8%, respectively, compared to control infected midguts (Figure 3A, Supplementary Figure 2D). This decrease in ISC proliferation was not due to reduced progenitor number caused by *licorne* depletion with *esg^ts^* and *P.e.* infection as the mean percent *esg^+^* progenitors in these midguts was similar to uninfected and *P.e*- infected control midguts (Supplementary Figure 2B-C). These data suggest that Licorne, likely through p38a and p38b, promotes ISC proliferation after *P.e.* infection.

**Figure 3.**
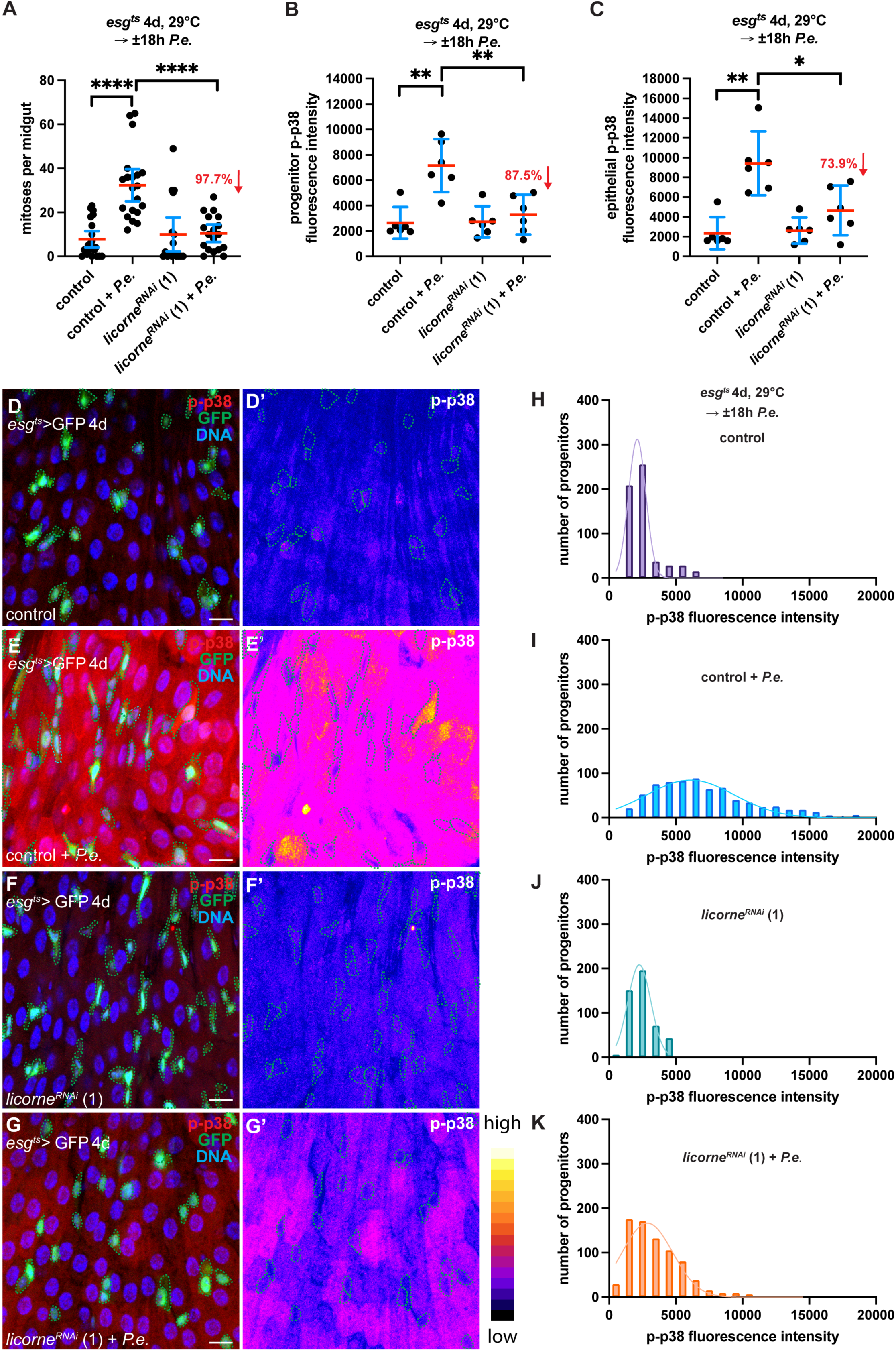
MKK3/Licorne is required in progenitors for ISC proliferation and p38 activation throughout the midgut epithelium after pathogenic bacterial infection. Licorne is required in progenitors for ISC proliferation after *P.e.* infection **(A).** Depleting *licorne* by expressing *licorne ^RNAi^* (1) in progenitors for 4 days with *esg^ts^* suppresses ISC proliferation by 97.7% compared to control *P.e.*-infected midguts (Mann–Whitney; p < 0.0001) (A). The mean number of PH3-positive cells per midgut with mean and 95% CI are shown for control midguts (n = 21 from one experiment), *P.e.*-infected control midguts (n = 20 from one experiment), midguts expressing *licorne ^RNAi^* (1) in progenitor cells for 4 days (n = 17 from one experiment) and *P.e.-* infected midguts expressing *licorne ^RNAi^* (1) in progenitors for 4 days (n = 18 from one experiment) (A). In A, the mean number of mitoses per midgut increased in *P.e.-*infected control midguts compared to uninfected control midguts (Mann-Whitney; p<0.0001). Licorne is required in progenitors for p38 activation in progenitors and throughout the midgut epithelium after *P.e.* infection **(B-K)**. p38 activation is blocked in progenitors and throughout the midgut epithelium of *P.e.*-infected midguts expressing *licorne ^RNAi^* (1) in progenitors by 87.5% (Mann-Whitney; p-value = 0.0087) and 73.9% (Mann-Whitney; p-value = 0.0260), respectively, compared to control *P.e.*-infected midguts (B-C). In B, the mean p-p38 fluorescent intensity in progenitors per midgut with mean and 95% CI are shown for control midguts (n = 6 from one experiment), *P.e.*-infected control midguts (n = 6 from one experiment), midguts expressing *licorne ^RNAi^* (1) in progenitors for 4 days (n = 6 from one experiment) and *P.e.*-infected midguts expressing *licorne ^RNAi^* (1) in progenitors for 4 days (n = 6 from one experiment). In C, the epithelial p- p38 fluorescence intensity per midgut (from those analysed in B) are shown with mean and 95%CI. In B and C, the mean p-p38 fluorescence intensity increased in *P.e.-*infected control midguts compared to uninfected control midguts (Mann-Whitney; p value = 0.0043 in B; Mann-Whitney; p-value = 0.0022 in C). p-p38 (D, E: red; D’, E’: LUT) increases in progenitors (E: GFP; E-E’: green-dashed outline) and throughout the midgut epithelium after *P.e.* infection compared to control (D-D’). Depleting *licorne* from progenitors for 5 days blocks the increase in p-p38 in progenitors and throughout the midgut epithelium (G-G’) compared to *P.e*.-infected control midguts (E-E’). The distribution (histogram) of p-p38 fluorescence intensity per progenitor is shown for uninfected control midguts (H), *P.e.-* infected control midguts (I), midguts expressing *licorne^RNAi^* in progenitors (J) and *P.e.-* infected midguts expressing *licorne^RNAi^* in progenitors (K)(progenitors from B). DNA is in blue. Scale bars are 20μm.

We next examined whether MKK3/Licorne activity in progenitors regulated p38 activation in both the progenitors and throughout the midgut epithelium. We indeed found that depletion of *licorne* with *esg^ts^* decreased p38 activation in both the progenitors and throughout the midgut epithelium (Figure 3B-C, Supplementary Figure 2E-F). Depleting *licorne* in progenitors with two RNAis (RNAis (1) and (2)) decreased p38 activation within progenitors by 87.5% and 93.8%, respectively, after *P.e.* infection compared to progenitors in control midguts (Figure 3B, 3D-K, Supplementary Figure 2E). Further, the same depletion of *licorne* in progenitors decreased p38 activation throughout the midgut epithelium by 73.9% and 61.6% compared to control infected midguts (Figure 3C-G’, Supplementary Figure 2F). These data suggest that MKK3-p38 signalling in progenitors after *P.e.* infection stimulates ISC proliferation and activates p38 throughout the midgut epithelium.

### Peptidoglycan recognition by intestinal progenitors stimulates their proliferation via MKK3-p38 signalling after pathogenic bacterial infection

Although we found that MKK3/Licorne can activate p38 within progenitors and promote ISC proliferation after *P.e.* infection, it was unclear how MKK3-p38 signalling was triggered in progenitors upon infection. As midgut ECs have been shown to activate p38 signalling after recognition of *Ecc15* peptidoglycan through PGRP-LC ^33^, we speculated that MKK3-p38 signalling in progenitors might be induced by directly sensing pathogenic bacteria during infection.

To test whether p38 activation in progenitors after gram-negative bacterial infection requires sensing of Dap-type peptidoglycan, we depleted the peptidoglycan receptors PGRP-LC and PGRP-LE in progenitors. We found that both receptors were required for ISC proliferation after *P.e.* infection (Figure 4A, Supplementary Figure 3A and Figure 5A).

**Figure 4.**
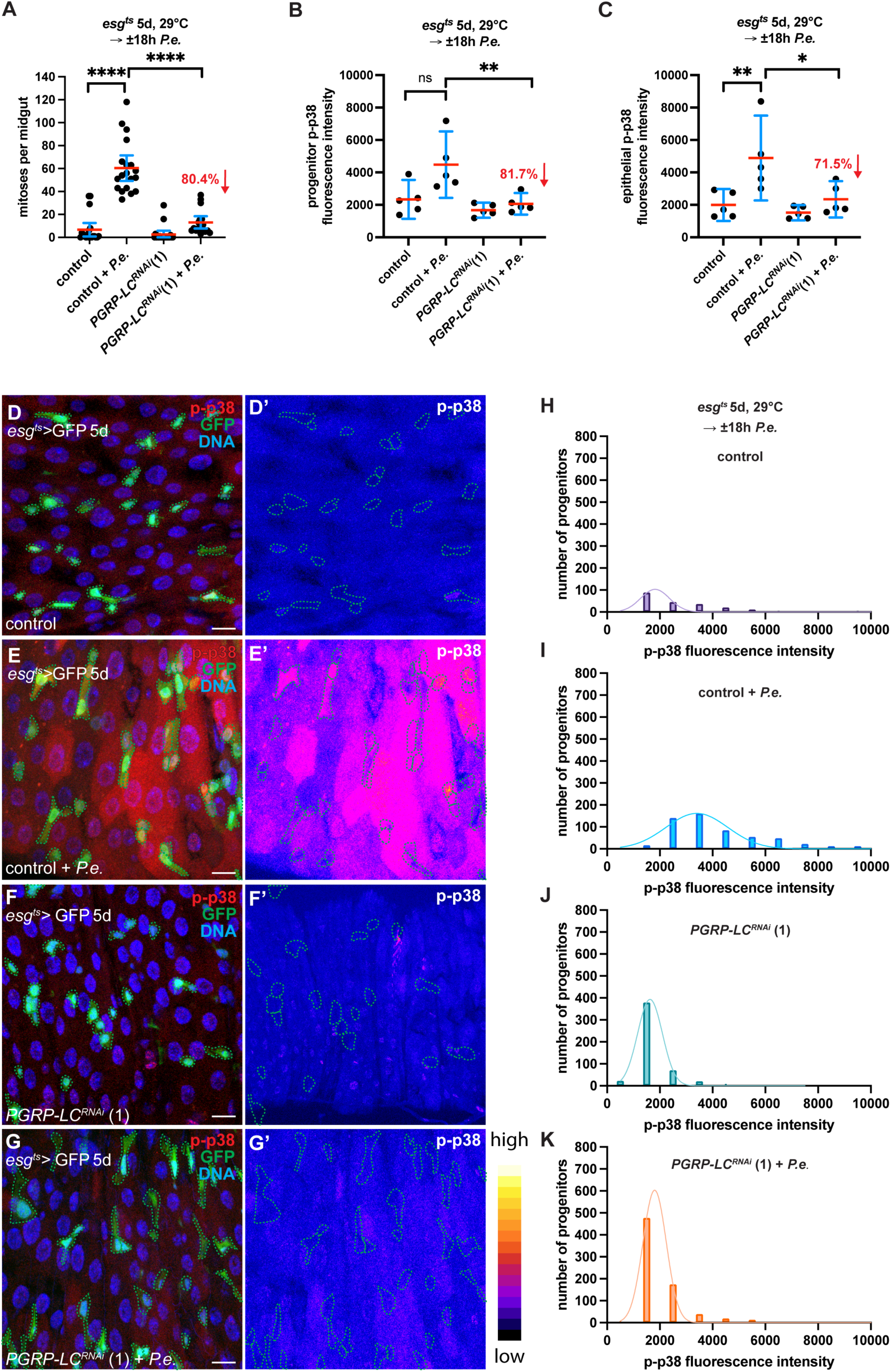
Peptidoglycan sensing by PGRP-LC is required in progenitors for ISC proliferation and p38 activation throughout the midgut epithelium after pathogenic bacterial infection. PGRP-LC is required in progenitors for ISC proliferation after *P.e.* infection **(A).** Depleting *PGRP-LC* by expressing *PGRP-LC ^RNAi^* (1) in progenitors for 5 days with *esg^ts^* suppresses ISC proliferation by 80.4% compared to control *P.e.*-infected midguts (Mann–Whitney; p < 0.0001) (A). The mean number of PH3-positive cells per midgut with mean and 95% CI are shown for control midguts (n = 19 from one of two experiments), *P.e.*-infected control midguts (n = 19 from one of two experiments), midguts expressing *PGRP-LC ^RNAi^* (1) in progenitor cells for 4 days (n = 20 from one of two experiments) and *P.e.-* infected midguts expressing *PGRP-LC ^RNAi^* (1) in progenitors for 4 days (n = 17 from one of two experiments) (A). In A, the mean number of mitoses per midgut increased in *P.e.-* infected control midguts compared to uninfected control midguts (Mann-Whitney; p<0.0001). PGRP-LC is required in progenitors for p38 activation in progenitors and throughout the midgut epithelium after *P.e.* infection **(B-K)**. p38 activation is blocked in progenitors and throughout the midgut epithelium of *P.e.*-infected midguts expressing *PGRP-LC ^RNAi^* (1) in progenitors by 81.7% (Mann-Whitney; p-value = 0.0079) and 71.5% (Mann-Whitney ; p-value = 0.0159), respectively, compared to control *P.e.*-infected midguts (B-C). In B, the mean p-p38 fluorescent intensity in progenitors per midgut with mean and 95% CI are shown for control midguts (n = 5 from one of two experiments), *P.e.*-infected control midguts (n = 5 from one of two experiments), midguts expressing *PGRP-LC ^RNAi^* (1) in progenitors for 5 days (n = 5 from one of two experiments) and *P.e.*-infected midguts expressing *PGRP-LC ^RNAi^* (1) in progenitors for 5 days (n = 5 from one of two experiments). In C, the epithelial p-p38 fluorescence intensity per midgut (from those analysed in B) are shown with mean and 95%CI. In B and C, the mean p-p38 fluorescence intensity increased in *P.e.-*infected control midguts compared to uninfected control midguts (Mann-Whitney; p value = 0.0556 in B; Mann-Whitney ; p-value = 0.0079 in C). p-p38 (D, E: red; D’, E’: LUT) increases in progenitors (E: GFP; E-E’: green-dashed outline) and throughout the midgut epithelium after *P.e.* infection compared to control (D-D’). Depleting *PGRP-LC* from progenitors for 5 days blocked the increase in p-p38 in progenitors and throughout the midgut epithelium (G-G’) compared to *P.e*.-infected control midguts (E-E’). The distribution (histogram) of p-p38 fluorescence intensity per progenitor is shown for uninfected control midguts (H), *P.e.-*infected control midguts (I), midguts expressing *PGRP-LC^RNAi^* (1) in progenitors (J) and *P.e.-*infected midguts expressing *PGRP-LC^RNAi^* (1) in progenitors (K) (progenitors from B). DNA is in blue. Scale bars are 20μm.

**Figure 5.**
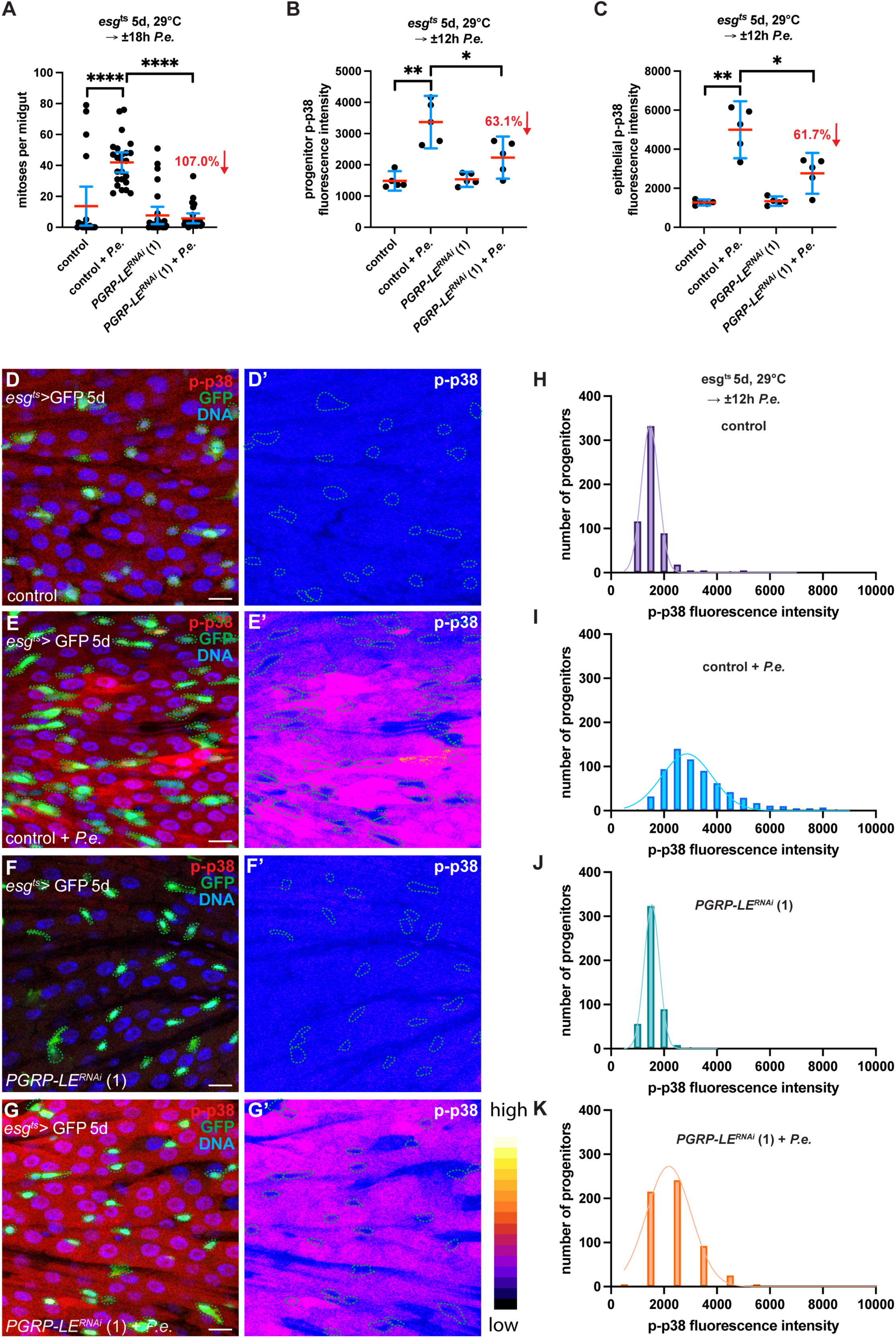
Peptidoglycan sensing by PGRP-LE is required in progenitors for ISC proliferation and p38 activation throughout the midgut epithelium after pathogenic bacterial infection. PGRP-LE is required in progenitors for ISC proliferation after *P.e.* infection **(A).** Depleting *PGRP-LE* by expressing *PGRP-LE ^RNAi^* (1) in progenitors for 5 days with *esg^ts^* suppresses ISC proliferation by 107.0% compared to control *P.e.*-infected midguts (Mann–Whitney; p < 0.0001) (A). The mean number of PH3-positive cells per midgut with mean and 95% CI are shown for control midguts (n = 20 from one of two experiments), *P.e.*-infected control midguts (n = 24 from one of two experiments), midguts expressing *PGRP-LE ^RNAi^* (1) in progenitor cells for 5 days (n = 28 from one of two experiments) and *P.e.-* infected midguts expressing *PGRP-LC ^RNAi^* (1) in progenitors for 5 days (n = 25 from one of two experiments) (A). In A, the mean number of mitoses per midgut increased in *P.e.-*infected control midguts compared to uninfected control midguts (Mann-Whitney; p<0.0001). PGRP-LE is required in progenitors for p38 activation in progenitors and throughout the midgut epithelium after *P.e.* infection **(B-K)**. p38 activation is blocked in progenitors and throughout the midgut epithelium of *P.e.*-infected midguts expressing *PGRP-LE ^RNAi^* (1) in progenitors by 63.1% (Mann–Whitney; p-value = 0.0317) and 61.7% (Mann–Whitney; p-value = 0.0317), respectively, compared to control *P.e.*-infected midguts (B-C). In B, the mean p-p38 fluorescent intensity in progenitors per midgut with mean and 95% CI are shown for control midguts (n = 5 from one of two experiments), *P.e.*-infected control midguts (n = 5 from one of two experiments), midguts expressing *PGRP-LE ^RNAi^* (1) in progenitors for 5 days (n = 5 from one of two experiments) and *P.e.*-infected midguts expressing *PGRP-LE ^RNAi^* (1) in progenitors for 5 days (n = 5 from one of two experiments). In C, the epithelial p-p38 fluorescence intensity per midgut (from those analysed in B) are shown with mean and 95%CI. In B and C, the mean p-p38 fluorescence intensity increased in *P.e.-*infected control midguts compared to uninfected control midguts (Mann–Whitney; p value = 0.0079 in B; Mann–Whitney; p-value = 0.0079 in C). p-p38 (D, E: red; D’, E’: LUT) increases in progenitors (E: GFP; E-E’: green-dashed outline) and throughout the midgut epithelium after *P.e.* infection compared to control (D-D’). Depleting *PGRP-LE* from progenitors for 5 days blocks the increase in p-p38 in progenitors and throughout the midgut epithelium (G-G’) compared to *P.e*.-infected control midguts (E-E’). The distribution (histogram) of p-p38 fluorescence intensity per progenitor is shown for uninfected control midguts (H), *P.e.-*infected control midguts (I), midguts expressing *PGRP-LE^RNAi^* (1) in progenitors (J) and *P.e.-*infected midguts expressing *PGRP-LE^RNAi^* (1) in progenitors (K)(progenitors from B). DNA is in blue. Scale bars are 20μm.

Depletion of PGRP-LC in progenitors with *esg^ts^* blocked ISC proliferation after *P.e.* infection by 80.4% and 49.0% with RNAi(1) and RNAi(2), respectively, compared to control infected midguts (Figure 4A, Supplementary Figure 3A). Similarly, we found that PGRP-LE depletion resulted in an 107.0% decrease in ISC proliferation after *P.e.* infection compared to control infected midguts (Figure 5A). Furthermore, the decrease in ISC proliferation due to PGRP-LC and PGRP-LE depletion in progenitors was not due to a decrease in progenitor number as the mean percent progenitors in these midguts was similar to uninfected and infected control midguts (Supplementary Figure 3B-C). Together, these data suggest that DAP-type peptidoglycan recognition through PGRP-LC and PGRP-LE in progenitors stimulates ISC proliferation after *P.e.* infection.

Since PGRP-LC, PGRP-LE and p38 were required to promote ISC proliferation after *P.e.* infection, we determined whether p38 activation within progenitors required pathogen recognition through PGRPs. We found that PGRP-LC depletion with *esg^ts^* blocked p38 activation in progenitors by 81.7% after *P.e.* infection compared to control infected midguts (Figure 4B, 4D-K). Similarly, PGRP-LE depletion with *esg^ts^* blocked p38 activation in progenitors by 63.1% after *P.e.* infection compared to control infected midguts (Figure 5B, 5D-K). These data suggest that pathogenic bacterial recognition by progenitors through PGRP-LC and PGRP-LE promotes ISC proliferation through activation of MKK3-p38 signaling after *P.e.* infection.

Because we had found that blocking p38a, p38b and MKK3/Licorne activity in progenitors decreased p38 activation throughout the midgut epithelium, we wondered whether PGRP-LC and PGRP-LE were required for this non-cell autonomous eXect as well. Indeed, depleting PGRP-LC and PGRP-LE in progenitors using *esg^ts^* blocked p38 activation throughout the midgut epithelium by 71.5% and 61.7%, respectively, after *P.e.* infection compared to control (Figure 4C, 4D-G’ and 5C, 5D-G’). These data suggest that p38 activation throughout the midgut epithelium requires pathogenic bacterial recognition by progenitors after *P.e.* infection.

### PGRP-LC/LE-MKK3-p38 activity in progenitors activates p38 in the midgut epithelium after pathogenic bacterial infection

So far, we had shown that depleting p38a and p38b, MKK3, PGRP-LC and PGRP-LE in midgut progenitors blocked p38 activation after *P.e.* infection not only within progenitors but also throughout the epithelium. Furthermore, we had found that increasing ISC proliferation through MKK3/Licorne overexpression led to p38 activation in nearby ECs. Thus, we asked whether it was increased ISC proliferation due to increased PGRP-LC/LE-MKK3-p38 signalling in progenitors that stimulated p38 activation throughout the midgut epithelium or whether it was specifically due to p38 signalling. To test this, we blocked ISC proliferation by depleting the *Drosophila* cdc25 phosphatase homolog, String, in progenitors with *esg^ts^*. This blocked ISC proliferation by 99.4% after *P.e.* infection compared to control infected midguts (Figure 6A). The decrease in ISC proliferation was not caused by reduced progenitor number due to *string* depletion with *esg^ts^* and *P.e.* infection as the mean percent *esg^+^* progenitors in these midguts was similar to uninfected and *P.e*-infected control midguts (Supplementary Figure 3D). Furthermore, blocking ISC proliferation did not block p38 activation, neither within progenitors (Figure 6B, 6D-K) nor throughout the epithelium (Figure 6C, Figure 6D-G’). These data suggest that pathogenic bacterial recognition via PGRP-LC/LE-MKK3-p38 signalling in progenitors promotes p38 activation in the epithelial regenerative niche.

**Figure 6.**
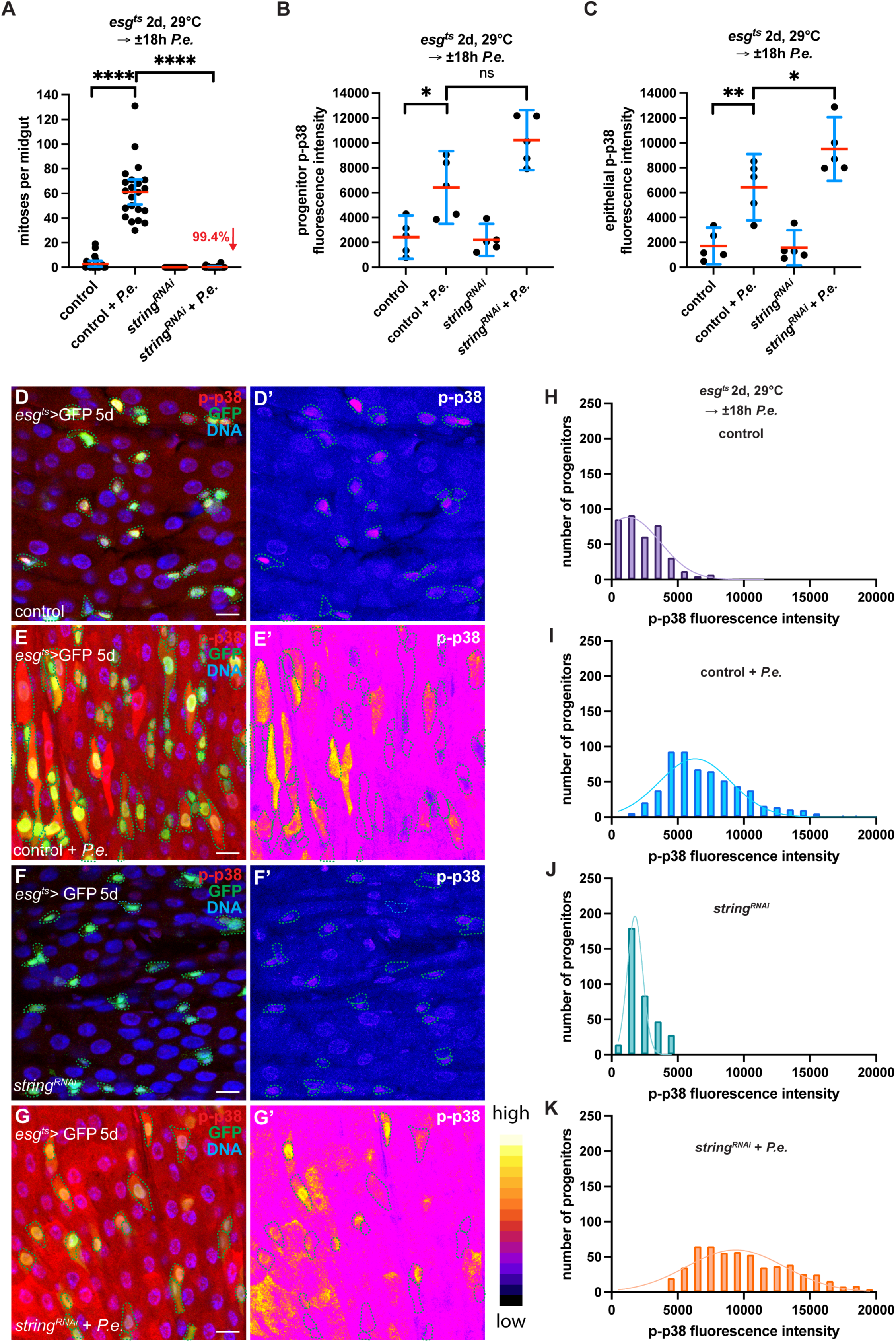
p38 activity in progenitors and not ISC proliferation induces p38 activation throughout the adult *Drosophila* midgut epithelium after pathogenic bacterial infection. ISC proliferation is not required for p38 activation in progenitors and throughout the midgut epithelium after *P.e.* infection **(A-C).** Depleting the cdc25-homologue *string* by expressing *string ^RNAi^* in progenitors for 2 days with *esg^ts^* suppresses ISC proliferation by 99.4% compared to control *P.e.*-infected midguts (Mann–Whitney; p < 0.0001) (A). The mean number of PH3-positive cells per midgut with mean and 95% CI are shown for control midguts (n = 22 from one experiment), *P.e.*-infected control midguts (n = 22 from one experiment), midguts expressing *string ^RNAi^* (1) in progenitors for 2 days (n = 25 from one experiment) and *P.e.-*infected midguts expressing *string ^RNAi^* (1) in progenitors for 2 days (n = 23 from one experiment) (A). In A, the mean number of mitoses per midgut increased in *P.e.-*infected control midguts compared to uninfected control midguts (Mann-Whitney; p<0.0001). ISC proliferation is not required for p38 activation in progenitors and throughout the midgut epithelium after *P.e.* infection **(B-K)**. p38 is activated in progenitors (Mann– Whitney; p-value = 0.0556) and throughout the midgut epithelium (Mann–Whitney; p-value = 0.0317) in *P.e.*-infected midguts expressing *string ^RNAi^* in progenitors compared to control *P.e.*-infected midguts (B-C). In B, the mean p-p38 fluorescent intensity in progenitors per midgut with mean and 95% CI are shown for control midguts (n = 5 from one of two experiments), *P.e.*-infected control midguts (n = 5 from one of two experiments), midguts expressing *string ^RNAi^* (1) in progenitors for 2 days (n = 5 from one of two experiments) and *P.e.*-infected midguts expressing *string ^RNAi^* (1) in progenitors for 2 days (n = 5 from one of two experiments). In C, the epithelial p-p38 fluorescence intensity per midgut (from those analysed in B) are shown with mean and 95%CI. In B and C, the mean p-p38 fluorescence intensity increased in *P.e.-*infected control midguts compared to uninfected control midguts (Mann–Whitney; p value = 0.0317 in B; Mann–Whitney; p-value = 0.0079 in C). p-p38 (D, E: red; D’, E’: LUT) increases in progenitors (E: GFP; E-E’: green-dashed outline) and throughout the midgut epithelium after *P.e.* infection compared to control (D-D’). p38 is activated in progenitors and throughout the midgut epithelium after *P.e.* infection despite blocking ISC proliferation by depleting *string* from progenitors for 2 days (G-G’). The distribution (histogram) of p-p38 fluorescence intensity per progenitor is shown for uninfected control midguts (H), *P.e.-*infected control midguts (I), midguts expressing *string^RNAi^* in progenitors (J) and *P.e.-*infected midguts expressing *string^RNAi^* in progenitors (K) (progenitors from B). DNA is in blue. Scale bars are 20μm.

In summary, we have uncovered a mechanism by which pathogenic bacterial infection promotes ISC proliferation in the adult *Drosophila* midgut. We find that DAP-type peptidoglycan sensing through PGRP-LC and PGRP-LE in progenitors, activates MKK3-p38 signalling within progenitors. This p38 activity stimulates ISC proliferation but also stimulates p38 activation throughout the midgut epithelium—a component of the regenerative niche (Figure 7).

**Figure 7.**
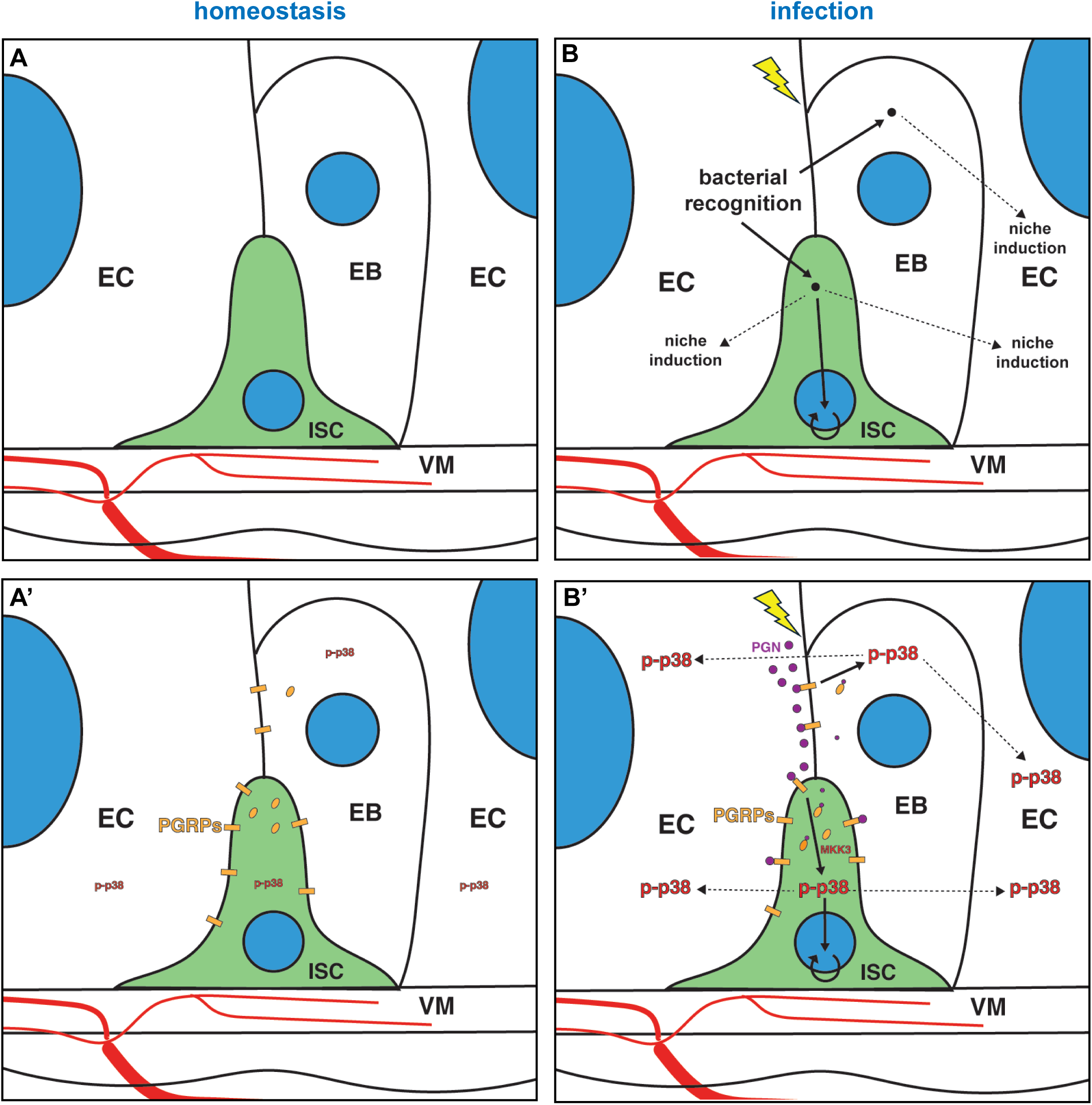
Recognition of pathogenic bacteria by intestinal progenitors via PGRP-MKK3-p38 signalling promotes adult *Drosophila* midgut regeneration. Under homeostasis, pathogenic bacteria are not recognised by adult *Drosophila* midgut progenitors (ISCs and EBs) and the levels of activated p-p38 remain low within progenitors and throughout the midgut epithelium **(A, A’)**. After infection, bacterial peptidoglycan is recognised by both ISCs and EBs, inducing MKK3-p38 signalling. In ISCs, MKK3-p38 signalling promotes ISC proliferation; in both ISCs and EBs, MKK3-p38 signalling stimulates p38 activation throughout the epithelial regenerative niche **(B, B’)**.

## Discussion

After infection by gram-negative pathogenic bacteria such as *Ecc15* and *P.e.,* p38 is known to be activated in the ECs of the adult fly midgut epithelium. Whilst *P.e.* infection activates both p38a and p38b via Nox-Ask1-MKK3 signalling, *Ecc15* infection induces p38a via PGRP- LC-IMD-Mekk1-MKK3 signalling in ECs ^33, 41^. Here we found that *P.e*. infection increases p38 activation in *esg^+^* midgut progenitors by 134.8% (Figure 1A-D). Furthermore, both p38a and p38b are required in these progenitors for ISC proliferation after infection as depletion of p38a and p38b in *esg^+^* progenitors reduced ISC proliferation by 73.3% (Figure 1E). Whilst the progenitor population includes ISCs, EBs and EEps, our data point to the requirement of p38 signalling mostly in ISCs. Targeted depletion of p38a and p38b specifically in ISCs with *esg^ts^Su(H)Gal80* reduced ISC proliferation by 59.6% (Figure 1F). This reduction is comparable to that observed in the broader progenitor population, suggesting that p38 signalling is primarily required in ISCs, rather than in EBs or EEps for ISC proliferation. In addition, we found that p38a and p38b were required in progenitors to activate p38 in a non-cell autonomous manner throughout the epithelium. Depletion of both p38a and p38b from progenitors reduced p38 activation throughout the midgut epithelium by 102.1% compared to control *P.e.*-infected midguts (Figure 1G). Together, these data suggest that p38 signalling in ISCs stimulates ISC proliferation and p38 activation throughout the midgut epithelium after pathogenic bacterial infection.

Since pathogen recognition by PGRP-LC can activate p38a signalling in ECs ^33^, we hypothesized that PGRPs may be required for p38 activation in ISCs to promote infection- induced proliferation. Indeed, PGRP-LC is highly expressed in midgut progenitors ^42^.

Further, general ISC proliferation is known to increase with age in *Drosophila*, and both PGRP-LC and PGRP-LE are required for this increase ^42^. We found that PGRP-LC and PGRP- LE were both required in progenitors for ISC proliferation after *P.e.* infection (Figure 4A and 5A) as depletion of either PGRP from progenitors strongly reduced ISC proliferation compared to control infected midguts (Figures 4A and 5A, Supplementary Figure 3A).

Further, depleting PGRP-LC and PGRP-LE with *esg^ts^* blocked p38 activation in progenitors after *P.e.* infection by 81.7% and 63.1%, respectively (Figures 4B and 5B). Together, these data suggest that recognition of pathogenic bacteria through PGRP-LC and PGRP-LE stimulates p38 signalling in progenitors and promotes ISC proliferation. Furthermore, our data suggest that pathogenic bacteria recognition by progenitors plays a significant role in the ISC response to *P.e.* infection and likely works synergistically with cues produced by the regenerative niche ^29^ that forms after midgut damage to stimulate full ISC proliferation.

Increased ISC proliferation after bacterial recognition may also serve as a strategy to repopulate the epithelium with uninfected cells.

Interestingly, overexpressing MKK3/Licorne to activate p38 in progenitor cells led to p38 activation not only within progenitors but also in neighbouring ECs (Figure 2A-B’).

Consistently, depletion of p38a and p38b, PGRP-LC and PGRP-LE in progenitors led to reduced p38 activation after infection not only within progenitors, but also throughout the epithelium (Figure 1G, 4B-C, 5B-C). Because p38 signalling controls ISC proliferation (Figure 1F), we wondered whether p38 activation throughout the epithelium was due to increased ISC proliferation. To test this, we blocked ISC proliferation by depleting the G2-M regulator, String, within progenitors prior to *P.e.* infection. Despite a 99.4% decrease in ISC proliferation after String depletion, p38 activation occurred throughout the epithelium after *P.e.* infection (Figure 6A, 6C). These data suggest that PGRP-LC/LE-p38 signalling in progenitors, and not ISC proliferation, promotes p38 activation throughout the midgut epithelium after *P.e.* infection. Furthermore, it suggests that pathogenic bacterial sensing in midgut progenitors can modify the regenerative niche to support regeneration. It is possible that p38 activity within progenitors stimulates p38 activation throughout the epithelium by a yet to be determined mechanism that influences Nox-Ask1 signalling in neighbouring ECs. Increased Nox-Ask1 signalling in ECs subsequently results in increased Upd3 expression that further supports ISC proliferation and midgut regeneration ^41^.

Previously, it was shown that damage sensing by ECs via Nox-Ask1 signalling activates p38 in ECs after *P.e.* infection ^41^. Our data suggest that p38 is activated in midgut ECs by pathogenic bacteria recognition by progenitors through PGRP-LC and PGRP-LE. For the case of *P.e.* infection, it is possible that both damage sensing in ECs and pathogen recognition in progenitors are important to activate p38 in ECs to suXiciently high levels.

Another study showed that p38a is activated in ECs after *Ecc15* infection by peptidoglycan recognition by PGRP-LC and IMD-Mekk1-MKK3 signalling ^33^. However, this study reduced MKK3 and PGRP-LC in the entire midgut epithelium, which could also reduce levels of MKK3 or PGRP-LC in progenitors. It therefore remains to be determined whether *Ecc15* infection is sensed by progenitors through the same pathway that we have identified here for *P.e.* infection.

The two PGRPs diXer in their peptidoglycan specificities: PGRP-LC recognises extracellular polymeric and monomeric DAP-type peptidoglycan, whereas PGRP-LE detects monomeric DAP-type peptidoglycan both extracellularly and intracellularly ^34, 36^. Our data suggest that progenitors sense both extracellular and intracellular peptidoglycan, since we find that both PGRPs are required in progenitors for ISC proliferation and p38 activation throughout the midgut epithelium. Intracellular detection of DAP-type peptidoglycan could occur through the entry of monomeric peptidoglycan or *P.e.* into midgut progenitors.

A similar mechanism may be in place in the mammalian intestine, with detection of bacterial peptidoglycan through Nod2 receptors influencing homeostatic epithelial growth. Nod2 sensing of the beneficial microbe *Lactiplantibacillus plantarum* has been shown to cause an increase in proliferative cell numbers in the small intestines of mice, alleviating stunted postnatal growth caused by undernutrition ^32^. Furthermore, Nod2 is highly expressed in Lgr5^+^ ISCs in intestinal organoids, and treatment of intestinal organoids with muramyl dipeptide (MDP), a synthetic peptidoglycan fragment, increased organoid growth and cell survival in response to stress ^31^. Our work provides an explanation for this result and indicates the potential for Nod2 in ISCs or intestinal progenitors to detect invading pathogenic bacteria and promote regenerative growth. Enteropathogenic *E. coli* (EPEC) is a major cause of diarrheal-related deaths worldwide, often due to dehydration and sepsis ^43^. Thus, identifying targets for therapies that stimulate the regenerative response would be beneficial to restore water absorption and barrier function. In support of a role for Nod2 receptors in intestinal regeneration, mutations in the Nod2 gene increase the risk of developing the inflammatory bowel disease, Crohn’s disease ^44^. Intestines of Crohn’s patients are severely damaged and fail to regenerate ^45^. Therefore, loss of bacterial detection by Nod2, particularly in ISCs, in inflammatory bowel diseases may contribute to reduced intestinal epithelial restitution in some Crohn’s disease patients. Treatments that enhance or restore bacterial recognition in intestinal progenitors may prove beneficial to these patients.

## Materials and Methods

### Fly stocks

The following fly stocks were used: w; *esgGAL4; tubGAL80^ts^ UAS-GFP* (*esg^ts^GFP*), *esgGAL4 UAS-2X EYFP; Su(H)Gbe-GAL80, tub-GAL80^ts^* (*esg^ts^ Su(H)GAL80*); *UAS-p38a+b^RNAi^* (1)(34238GD)^41^, *UAS-p38a+b^RNAi^* (2)(52277GD)^41^, *UAS-licorne^RNAi^* (1)(106822KK)^41^, *UAS- PGRP-LC^RNAi^* (1)(51968GD), *UAS-PGRP-LC^RNAi^* (2)(101636KK) ^42, 46^ and *UAS-PGRP-LE^RNAi^* (1)(108199KK) ^42^, KK control line (60100KK) from the Vienna *Drosophila* Stock Center (VDRC) and *UAS-mCherry^RNAi^*, *UAS-licorne^RNAi^* (2)(JF01433), *UAS-string^RNAi^* (HMC00146) from the Bloomington *Drosophila* Stock Center (BDSC); *UAS-licorne* ^41^ from the Zürich Fly ORFeome project (FlyORF) and *UAS-p38b^antisense^* ^47^ from the Kyoto Stock Center.

### Drosophila genetics

Flies raised at 18°C were shifted to 29°C to induce GAL4-mediated *UAS*-transgene expression. Experiments were performed using 5-10 day old, adult, female *Drosophila melanogaster*. Typically, between 10-20 flies were used per experiment. Flies were first selected based on genotype, then randomly chosen for use in experiments.

### Pathogenic bacterial infection

*Pseudomonas entomophila* (*P.e.*) infection: flies were exposed for 18 hours with food supplemented with 500ul of 5% sucrose or 20x *P.e.* resuspended in 5% sucrose, prepared from an 18-hour culture grown at 30°C, 130 RPM.

### Histology

Adult *Drosophila* midguts were dissected in cold PBS and then fixed in 8% formaldehyde containing phosSTOP (Roche) for 1 hour. After fixation, midguts were washed at least three times in PBS, 0.1% Triton X-100 for 10-15 minutes. Midguts were then blocked for at least 30 minutes in blocking solution containing PBS, 0.1% Triton X-100, 1% bovine serum albumin (BSA) and 2% normal goat serum (NGS). After blocking, midguts were incubated overnight with primary antibodies in blocking solution at 4°C. The following primary antibodies were used: rabbit polyclonal anti-phospho Ser 10 histone 3 (Millipore; 06-570; 1:1000), mouse monoclonal anti-phospho Ser 10 histone 3 (Sigma-Aldrich; 3H10; 05-806; 1:1000), rabbit monoclonal anti-phospho p38 (Thr180/Tyr182) (3D7)(Cell Signaling; 9215; 1:50) and rabbit polyclonal anti-phosphorylated p38 (Thr180/Tyr182) (Cell Signaling; 9211; 1:200). Midguts were then washed three times as above in PBS containing 0.1% Triton X-100, stained in PBS, 0.3% Triton X-100, 0.1% BSA with Alexa Fluor-conjugated secondary antibodies (Life Technologies, 1:1000) and Hoechst 33258 (Life Technologies) and mounted in Vectashield (Vector Laboratories).

### Microscopy

Samples were analysed using a Leica DM16000 and Leica SP5 microscope. Images were processed using ImageJ (NIH) and Adobe Photoshop 26.1.0. Confocal images are presented as maximum intensity projections of images obtained every 0.5-1 um. Each z- stack was acquired with the same laser intensity and gain.

The number of mitoses per midgut was determined by counting the number of phospho-Ser 10 histone 3-positive cells from whole adult female midguts from two or more independent experiments. The mean number of mitoses per midgut and 95% confidence interval are presented for each genotype and treatment.

### Image quantitation

*Phospho-p38 in progenitors:* sum intensity projections of 10 slices from 0.5 μm z-stacks of the progenitor layer were generated for the esg>GFP and phospho-p38 channels using ImageJ. *esg*>GFP progenitors were selected, and the ROIs were recorded. These ROIs were then applied onto sum intensity projections of the p-p38 channel to measure the raw integrated density within progenitors. The raw integrated density of each progenitor was divided by the area of each progenitor to obtain the normalised fluorescence intensity of p- p38 per progenitor.

*Phospho-p38 in the midgut epithelium:* sum intensity projections of 10 slices from 0.5 μm z-stacks of the progenitor layer for the p-p38 channel were generated using imageJ. The freehand selection tool was used to trace around the epithelium to measure the raw integrated density and epithelial area. The raw integrated density of p-p38 was then divided by the area of selected epithelium to obtain the normalised fluorescence intensity of p-p38 in the whole epithelium.

*% esg-positive cells:* maximum intensity projections of 10 slices from the progenitor layer from 0.5 or 1 μm z-stacks were generated for the esg>GFP and DNA (Hoechst) channels in ImageJ. The total number of *esg^+^* cells with small nuclei and the total number of nuclei were counted. The number of esg^+^ cells was then divided by the total number of nuclei to obtain % *esg-*positive cells.

### Statistical analysis

Statistical analyses were conducted using GraphPad Prism 10. A two-sided Mann-Whitney test or an unpaired t-test was applied to determine statistical significance. The significance level is denoted by an * for p<0.05, ** for p<0.01, *** p<0.001, and **** for p<0.0001, and ns, for not significant, p>0.05.

## Acknowledgements

We thank the Bloomington *Drosophila* Stock Center, Vienna *Drosophila* RNAi Center, Zürich Fly ORFeome project (FlyORF) and Kyoto Stock Center for fly stocks. This research was funded in whole by the Wellcome Trust and Royal Society (Grant Number 220198/Z/20/Z). For the purpose of open access, the author has applied a CC BY public copyright license to any Author Accepted Manuscript version arising from this submission.

## Author contributions

P.H.P. conceived, developed and supervised the project. P.H.P. designed the experiments. B.U., R.W. and S.C. performed and analysed the experiments. B.U. contributed Figures 1G, Figures 3-6, Supplementary Figure 1B-G, Supplementary Figure 2B-F and Supplementary Figure 3; R.W. contributed Figures 1E-F, Figure 2D and Supplementary Figure 1A; S.C. contributed Figure 1A-D, Figure 2A-C and Supplementary Figure 2A. B.U. and P.H.P prepared the figures and figure legends. P.H.P. wrote and edited the manuscript.

## Conflict of Interests

The authors declare they have no conflict of interest.

